# Preserved Intrinsic Neural Timescale Organization with Hierarchical Variation in Autism Spectrum Disorder

**DOI:** 10.64898/2026.02.27.708484

**Authors:** Yumi Shikauchi, Ryuta Aoki, Takashi Itahashi, Masaaki Shimizu, Taiga Naoe, Tsukasa Okimura, Haruhisa Ohta, Ryu-ichiro Hashimoto, Motoaki Nakamura

## Abstract

Intrinsic neural timescales (INTs) index the temporal decay of neural activity and form a cortical hierarchy from fast sensorimotor to slow transmodal regions. Altered INTs have been reported in autism spectrum disorder (ASD), but it remains unclear whether the hierarchical organization is preserved and how individual variability along this hierarchy relates to sensory traits. Using resting-state fMRI from 182 participants (67 ASD, 115 typically developed controls (TDC)), we estimated INT at each cortical vertex from the autocorrelation half-life and averaged these values across four five-minute runs per participant. Vertex-wise INTs were then averaged within predefined cortical parcels and large-scale functional networks for subsequent analyses.

The cortical INT hierarchy was preserved in ASD, showing comparable sensorimotor-to-transmodal hierarchy in both groups. However, regions operating at longer timescales showed prolonged INTs in ASD, and such tendency increased systematically along the hierarchy. No vertex, parcel, or network survived correction of multiple comparisons, indicating that observed alterations followed a distributed hierarchical trend rather than a focal pattern. To disentangle group-level differences from inter-individual variability, we next modeled each participant’s parcel-wise INT profile relative to a TDC-derived group-averaged template. At the individual level, decomposition of INT profiles revealed that global shifts and hierarchical scaling primarily reflected demographic variation (plimarily sex) rather than diagnostic group membership. After accounting for these components, residual deviations from theTDC-derived cortical INT hierarchy showed a modest association with sensory traits characterized by reduced sensory registration.

Together, these findings indicate that while the large-scale hierarchical organization of cortical temporal dynamics is largely preserved in ASD, individual-specific deviations from this hierarchy may contribute to variability in sensory experience beyond group-level differences.

**Highlights:**

1. Across both autism spectrum disorder (ASD) and typically developed controls (TDC), intrinsic neural timescales (INTs) followed the established sensory cortical hierarchy and showed a negative association with cortical microstructural markers (myelin content and neurite density index), with no group difference in hierarchical slope.
2. At the whole-brain level, the hierarchical organization of INTs was preserved across ASD and TDC, although regions operating at longer timescales exhibited relatively greater extension in ASD.
3. Global shifts and hierarchical scaling, describing individual positioning within the INT-based cortical hierarchy, were more strongly associated with demographic variation (primarily sex) than with diagnosis.
4. After accounting for these global and hierarchical components, residual deviations from the INT-based cortical hierarchy were modestly associated with sensory traits characterized by reduced sensory registration.

## Introduction

The human brain processes information across multiple temporal scales, forming a hierarchical organization that mirrors its anatomical and functional architecture (Golesorkhi et al., 2021; Honey et al., 2012; Ito et al., 2020). The intrinsic neural timescale (INT) has been proposed as one of several indices to describe temporal hierarchy, indicating the duration over which neural activity remains autocorrelated within a region (Fig. 1a)(Murray et al., 2014). Neural activity in primary sensorimotor areas fluctuates rapidly, reflecting short INT, whereas association cortices integrate information over longer timescales, supporting abstraction, prediction, and self-referential processing. This gradient of temporal integration, observed across sensorimotor-to-association hierarchies and conserved across mammalian cortices (Murray et al., 2014; Runyan et al., 2017), is considered a fundamental principle of cortical computation that enables the coordination of perception and cognition over time.

**Figure 1.**
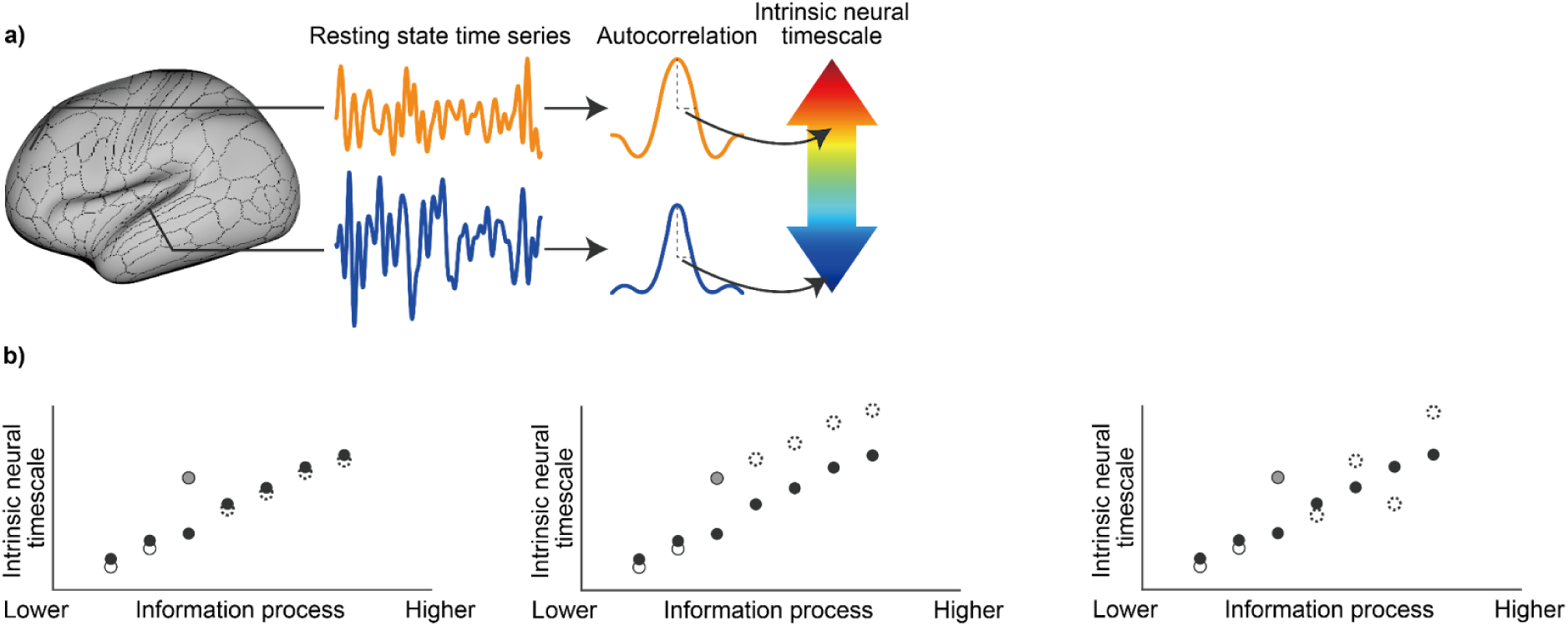
Conceptual illustration of the cortical hierarchy of Intrinsic neural timescales (INTs) and its individual-level deviations. (a) INT was computed from resting-state fMRI time series as half of the full width at half maximum of the autocorrelation function, representing the characteristic temporal span over which spontaneous neural activity remains correlated with itself. Regions with longer INTs (orange) exhibit more slowly decaying autocorrelation, whereas regions with shorter INTs (blue) show faster signal fluctuations. INT provides a quantitative summary of the temporal persistence of neural activity at each cortical region and is commonly used to characterize large-scale temporal hierarchies across the cortex. (b) Solid black circles represent the typical INT profile, reflecting the orderly increase of timescales along the cortical processing hierarchy. Open circles indicate the INT values of a given individual. A gray dashed circle highlights a local deviation at a specific region. Three possible patterns of INT modulation are illustrated. Left: a focal deviation confined to a single region, with no systematic propagation to other hierarchical levels. Center: a cascading deviation in which downstream regions along the information-processing hierarchy continue to shift in the same direction, preserving hierarchical order but altering its global scaling. Right: a destabilized pattern in which downstream regions exhibit irregular or non-monotonic deviations, suggesting disruption of hierarchical organization.

Individuals with autism spectrum disorder (ASD) exhibit atypical sensory experiences and difficulties in integrating and predicting information over time, suggesting alterations in both temporal and hierarchical organization of neural processing. Prior neuroimaging studies have reported regional alterations of INTs in ASD (Uscătescu et al., 2023; Watanabe et al., 2019), yet the reported effects have varied considerably in both direction and anatomical location. Such inconsistencies suggest that ASD-related differences in INTs may not be adequately described by region-wise comparisons alone, as these approaches do not account for inter-individual variation in glocal INT baselines or hierarchical scaling across the cortex. Consequently, local deviations may reflect shifts in the relative positioning of regions within the cortical hierarchy rather than isolated abnormalities (Fig. 1b). This perspective motivated the hypothesis that ASD involves system-level reorganization of cortical timescale hierarchy, rather than focal disruptions in specific regions (Bernhardt et al., 2025).

Gradient-based and network-level analyses indicate a reduced differentiation between sensorimotor and transmodal systems, suggesting that ASD may involve broad hierarchical modulation rather than focal dysfunction. This hierarchical imbalance could provide a unifying framework for understanding both enhanced perceptual sensitivity and difficulties in higher-order integration observed in ASD.

From a theoretical perspective, alterations in INT are unlikely to remain confined to isolated cortical loci (Chaudhuri et al., 2015; Murray et al., 2014). Because each region’s temporal integration depends on interactions with other levels of the cortical hierarchy, a local perturbation of timescales would disrupt the coordinated flow of information across the system. Therefore, deviations in INT are expected to be accompanied by distributed, hierarchical adjustments that preserve large-scale coherence in neural processing. Building on this framework, the present study aimed to examine whether ASD involves alterations in the large-scale organization of INT rather than focal abnormalities restricted to specific cortical regions. We analyzed resting-state fMRI data from a large cohort collected through the Brain/MINDS Beyond project (Koike et al., 2021), allowing for a comprehensive assessment of the cortical hierarchy in both ASD and typically developed controls (TDC). In addition, by relating INT to individual differences in sensory processing profiles, we sought to determine whether the association between neural timescales and sensory traits differs across diagnostic groups.

Based on these considerations, we examined whether ASD-related alterations in INT are better characterized as local changes confined to specific sensorimotor regions or as more globally distributed modulations across the cortical hierarchy. To place these temporal hierarchy patterns in a broader biological context, we also compared INT with microstructural indices, including T1w/T2w myelin maps (Glasser & Van Essen, 2011) and neurite density index (NDI) derived from diffusion-weighted imaging (Fukutomi et al., 2018; Zhang et al., 2012). These multimodal comparisons allow us to examine whether the large-scale organization of temporal processing aligns with, or departs from, underlying structural gradients. Furthermore, by examining how individual variability along the cortical INT hierarchy relates to sensory processing profiles, we sought to clarify whether deviations from large-scale temporal organization are associated with sensory atypicalities in ASD.

## Materials and Methods

### Participants

Data were collected as a part of Brain/MINDS Beyond projects (Koike et al., 2021), which established a high-quality scanning protocol, the Harmonization Protocol (HARP) to acquire multimodal brain images including T1- and T2-weighted, resting-state and task-based functional, and diffusion-weighted MRI. The present study analyzed resting-state data from 316 participants who met all inclusion criteria as follows. To minimize potential disparities introduced by variations in imaging equipment, only data acquired using the Skyra fit scanner were included. Participants were required to be labeled as either typically developed controls (TDC) or individuals with ASD, to have completed four resting-state fMRI runs, and to have fully responded to the Adolescent/Adult Sensory Profile (AASP)(Brown et al., 2001). Participants were excluded if they had any structural brain anomalies, were younger than 18 years, exhibited severe motion or imaging artifacts, had mean framewise displacement (FD) exceeding three standard deviations from the overall mean (FD > 0.178 mm). A clinical diagnosis of ASD was required for inclusion in the ASD group; individuals with neurodevelopmental diagnoses in the absence of ASD (e.g., Attention-Deficit/Hyperactivity Disorder (ADHD)-only) were excluded. Other psychiatric comorbidities were not systematically screened and therefore were not used as exclusion criteria. All participants were community-dwelling individuals capable of independently volunteering for MRI research. After applying these criteria, 115 TDC and 67 ASD participants remained for analysis (Table 1). Information regarding current medication use was collected. In the ASD group, three participants were taking methylphenidate (Concerta) and one was taking atomoxetine (Strattera), whereas no participants in the TDC group were receiving ADHD medication. Medication use was not used as an exclusion criterion. Estimated intelligence quotient (IQ) based on the Japanese adult reading test (JART) was reported for both groups. No TDC participants had JART scores below 85. The Wechsler Adult Intelligence Scale Fourth Edition (WAIS-IV) or Third Edition (WAIS-III) was administered in the ASD group. No ASD participants had both JART and WAIS-IV/WAIS-III scores below 85. IQ was therefore not used as an exclusion criterion. This study was approved by the institutional ethics committee of Showa Medical University (approval number: B-2014-019). All the participants provided written informed consent. All procedures were conducted according to the Declaration of Helsinki.

**Table 1.** Demographics of the analyzed population. IQ, intelligence quotient; JART: Japanese adult reading test; WAIS, Wechsler Adult Intelligence Scale.

| | # of participants | Age mean ± SD (min–max) | Frame-wise displacement mean ± SD (min–max) | Estimated IQ by JART mean ± SD (min–max) | IQ by WAIS IV mean ± SD (min–max) | IQ by WAIS III mean ± SD (min–max) |
| --- | --- | --- | --- | --- | --- | --- |
| ASD | 67<br>(Male 46, 69%) | 27.925 ± 7.768<br>(18–48) | 0.072 ± 0.027<br>(0.033–0.147) | 105.653 ± 7.743<br>(75.338–119.800) | 102.609 ± 14.693<br>(75–131)<br>(n = 41) | 101.800 ± 11.020<br>(78–118)<br>(n = 15) |
| TDC | 115<br>(Male 88, 77%) | 26.321 ± 6.375<br>(18–45) | 0.063 ± 0.024<br>(0.025–0.148) | 108.253 ± 7.743<br>(85.443–119.800) | | |

### MRI data acquisition

All MRI data were acquired following the HARP implemented in the Brain/MINDS Beyond project (Koike et al., 2021). Scanning was performed on a 3T Siemens MAGNETOM Skyra fit scanner equipped with a 32-channel head coil.

Structural T1-weighted and T2-weighted images were obtained using a high-resolution 3D MPRAGE and 3D SPACE sequences (0.8-mm isotropic), respectively. Diffusion-weighted images were acquired using a spin-echo echo planar imaging (EPI) sequence with 1.7-mm isotropic resolution and a multishell diffusion scheme (b = 0, 700, and 2000 s/mm²). Diffusion weighting was applied along multiple gradient directions for each shell, with phase-encoding directions alternated between anterior-to-posterior and posterior-to-anterior across acquisitions. Resting-state fMRI data were collected using a multiband EPI sequence (TR = 800 ms; 2-mm isotropic). Each participant completed four runs of resting-state fMRI (5 min per run), with two runs acquired in the anterior-to-posterior direction and two in the posterior-to-anterior direction.

### MRI data preprocessing

Preprocessing of the HARP data was performed using the Human Connectome Project (HCP) minimal preprocessing pipeline (Glasser et al., 2013). Briefly, this pipeline includes gradient distortion correction, motion correction, and EPI susceptibility-distortion correction using paired spin-echo EPI images with opposite phase-encoding directions, followed by registration to structural images and standard surface/volume spaces. Residual artifacts were further reduced using ICA-FIX denoising, applied in a multi-run manner to leverage the four resting-state runs per participant. The preprocessed fMRI data were then used for the estimation of INT maps in 32k Conte69 mesh surface space with native spatial resolution.

The T1w/T2w (myelin) maps (Glasser & Van Essen, 2011) in 32k Conte69 mesh surface space were used to compare functional (INT) and structural (T1w/T2w) measures of hierarchy. In addition, NDI maps derived from diffusion-weighted imaging were employed to assess microstructural hierarchy. Following Fukutomi et al. (2018), diffusion-weighted signals were fitted with a multi-compartment model, and the volume fraction of the intra-cellular compartment was used as NDI (Fukutomi et al., 2018).These maps were also resampled to the 32k Conte69 mesh surface space for comparison with the functional and myelin measures.

### INT Calculation

Before estimating the vertexwise INT values, preprocessed fMRI data were further processed with the following steps: (1) regression of nuisance variables, including the top aCompCor components (five components each from white-matter and cerebrospinal-fluid masks), global-brain signal, and the 24 motion parameters along with their first derivatives, resulting in a total of 30 nuisance regressors (Behzadi et al., 2007; Itahashi et al., 2025); (2) bandpass filtering in the 0.01–0.1 Hz range; (3) excluding volumes with FD greater than 0.5 mm, as well as the volumes immediately preceding and following those volumes, from subsequent analyses (Power et al., 2012).

The calculation of the INT was conducted as proposed in a previous study (Raut et al., 2020). Briefly, for each cortical region, we computed the autocorrelation function (ACF) of the resting-state time series and quantified the time lag at which the ACF decayed to half of its maximum, reflecting the temporal persistence of neural activity. The vertexwise INT values were then averaged across the four runs to obtain a stable estimate for each participant. To obtain parcel-and network-level measures, run-averaged vertexwise INT values were aggregated within Glasser’s multimodal cortical 360 parcels and subsequently within the 12 functional networks defined by the Cole–Anticevic Brain-wide Network Partition (Glasser et al., 2016; Ji et al., 2019).

### Structure–function correspondence analysis

To assess structure–function correspondence within sensory hierarchies, we computed spatial correlations between regional INTs and microstructural measures (myelin content and NDI). For each participant and within each sensory system, Spearman’s rank correlations were calculated across regions constituting the predefined histological hierarchy (Wengler et al., 2020), controlling for age, sex, and mean FD. The resulting participant-level correlation coefficients were then compared between groups using ANCOVA, with age, sex, and FD included as covariates.

### Quantification of individual deviations relative to the empirical hierarchical ordering of INTs

To evaluate individual variability in INTs beyond global shifts and hierarchical scaling, we characterized subject-specific deviations relative to an empirically derived INT hierarchy.

To establish a reference for this canonical organization, an INT template was constructed by averaging parcel-wise INT values across TDC. This template defined an INT-based cortical hierarchy, against which individual INT profiles were evaluated.

For each participant *i*, parcel-wise INT values *y_i,p_* were modeled as a linear transformation of the template INT values *t_p_* using ordinary least squares regression:

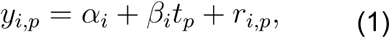

where *α_i_* captures a subject-specific global offset in INTs, *β_i_* reflects the degree to which the individual INT profile follows the hierarchical gradient of the template, and *r_i_*, represents residual deviations not explained by linear alignment to the template. This formulation allows separation of overall INT prolongation or shortening and hierarchical scaling effects from non-linear, spatially distributed deviations. To avoid circularity, when computing *t_p_* for participants in the TDC group, each participant’s own data was excluded from the TDC reference distribution used to derive the hierarchy.

Individual variability in INT distribution was quantified as the root mean square (RMS) of the residuals across all parcels:

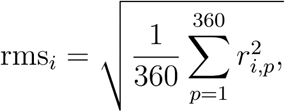

where *r_i,p_* were computed after linear alignment of individual INT profiles to the TDC-derived template. As the analysis was performed on a fixed parcellation comprising 360 cortical parcels, this metric provides a consistent measure of distributed deviations from the INT-based cortical hierarchy across participants.

To relate individual INT deviations to sensory traits, we analyzed associations between rms and sensory processing measures assessed by AASP. Given substantial correlations among the four AASP subscale scores (Turjeman-Levi & Kluger, 2022), principal component analysis (PCA) was applied to derive orthogonal sensory dimensions. To assess relationships between individual INT deviations and sensory traits, we computed Spearman’s rank partial correlations between the residual RMS measure of INT and AASP principal component (PC) scores, while controlling for age, sex, and FD.

## Results

### INTs reflect sensory processing hierarchies and align with microstructural markers

We estimated INTs from resting-state fMRI by quantifying the half width at half maximum of the autocorrelation function for each cortical region (Fig. 1a)(Ito et al., 2020; Raut et al., 2020). To confirm whether the previously reported hierarchical organization of INTs was also present in individuals with ASD, we checked the hierarchical gradient of INT within the auditory, visual, and somatosensory processing regions. These sensory systems were selected because their hierarchical organization—from primary to higher-order association areas—has been systematically delineated in previous anatomical and functional studies, providing a well-defined scaffold against which the cortical hierarchy of INTs can be evaluated.

Within auditory processing regions, INTs exhibited significant correlations with the a priori defined anatomical hierarchy described by Wengler et al. (2020)(Fig. 2a). The Spearman correlation between participant-level INT and the histological hierarchy was significant in both groups (ASD: *r_s_*= 0.808, *p* < 1 × 10^-5^; TDC: *r_s_* = 0.859, *p* < 1 × 10^-5^, after adjusting for age, sex, and FD). None of the covariates (age, sex, FD) showed significant effects (all *p* > 0.400). No significant group difference was observed in the participant-level correlation coefficients (ANCOVA including age, sex, and FD as covariates, *F*(4, 177) = 0.804, *p* = 0.523). The main effect of group did not reach statistical significance (*t*(177) = 1.415, *p* = 0.158). The effect size was small (Cohen’s d ≈ 0.21), indicating that the temporal hierarchy of INT within the auditory cortex was highly consistent across individuals, irrespective of diagnostic groups. This suggests that the basic organization of temporal processing in the auditory system is preserved in ASD, with only minimal deviation, if any.

**Figure 2.**
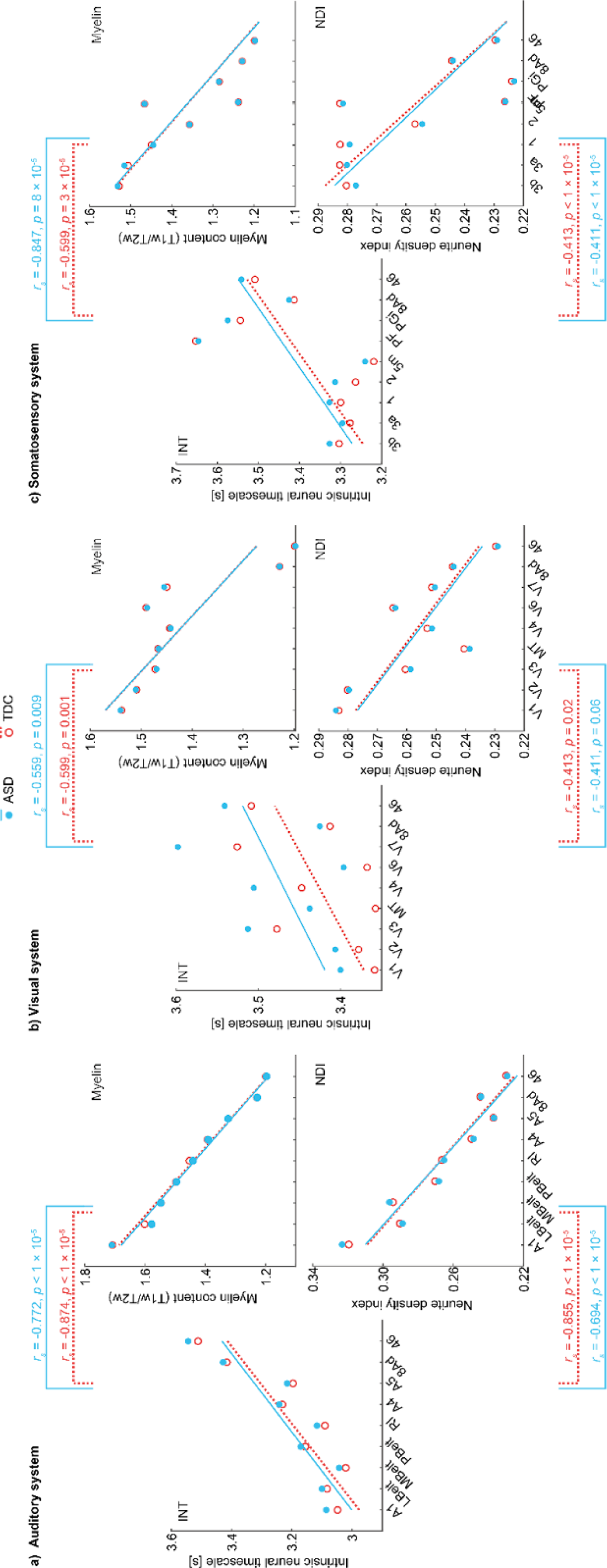
Cortical intrinsic neural timescale (INT) hierarchy across sensory systems and its association with microstructural profiles. (a) In both ASD (cyan) and TDC (red) groups, INTs exhibited hierarchical orderings that broadly aligned with the anatomically defined auditory (a), visual (b), and somatosensory (c) hierarchies reported by Wengler et al. (2020). While the overall monotonic trend was preserved across modalities, the strength of the gradient varied, with less uniform ordering observed in the somatosensory hierarchy, consistent with prior reports. Each circle represents an estimated marginal mean adjusted for age, sex, and frame-wise displacement, and partial correlations were computed with the same covariates. At the participant level, the spatial profiles of INTs across hierarchical levels was significantly correlated with the corresponding microstructural profile with each group. Specifically, the regional distribution of INT values showed negative spatial correspondence with both myelin content and neurite density index, indicating that regions characterized by denser microstructure tend to exhibit shorter intrinsic timescales.

Similar results were observed in the visual and somatosensory processing hierarchies (Fig. 2b, c). In the visual system, participant-level INT profiles were significantly correlated with the predefined histological hierarchy in both groups (ASD: *r_s_* = 0.539, *p* = 0.017; TDC: *r_s_*= 0.567, *p* = 0.004) with no significant group differences (*p* = 0.307). In the somatosensory processing system, INTs also showed significant correspondence with the histological hierarchy in both groups (ASD: *r_s_* = 0.744, *p* = 2 × 10^-4^; TDC: *r_s_* = 0.599, *p* = 3 × 10^-4^) with no significant group differences (*p* = 0.354). Although the hierarchical gradient appeared less strictly monotonic compared to auditory and visual systems—consistent with prior reports (Wengler et al., 2020). Together, these findings demonstrate that the hierarchical ordering of INTs within sensory systems is largely comparable between ASD and TDC.

To further validate the reliability of the temporal hierarchy captured by INT in individuals with ASD, we compared it with structural measures known to reflect cortical hierarchy. Specifically, we examined the relationships between INT, cortical myelin content, and NDI, both of which have been established as structural proxies of hierarchical organization (Burt et al., 2018; Fukutomi et al., 2018; Huntenburg et al., 2018; Paquola et al., 2019). Within each individual, we computed individual-level pairwise correlations between regional INTs and myelin content, as well as between INTs and NDI. Both groups showed significant negative correlations between INT and myelin content (ASD: *r_s_* =-0.772, *p* < 1 × 10^-5^; TDC: *r_s_* =-0.874, *p* < 1 × 10^-5^) and between INT and NDI (ASD: *r_s_* =-0.694, *p* < 1 × 10^-5^; TDC: *r_s_* =-0.855, *p* < 1 × 10^-5^), after controlling for age, sex, and FD (Fig. 1b). These results replicate the previously reported negative relationship between INTs and myelin density in healthy adults (Ito et al., 2020) and confirm that the hierarchical gradient of INT is tightly coupled with structural hierarchy in both groups.

Similar structure–function correspondence was observed in the visual and somatosensory systems. In the visual hierarchy, INT profiles showed significant negative spatial correlations with myelin content in both groups (all *p* < 0.05). Negative associations with NDI were also observed in both groups, although the magnitude of these correlations was comparatively modest (Fig. 2b). In the somatosensory system, INT demonstrated significant negative correlations with both myelin content and NDI in both groups (all *p* < 1 × 10^-4^)(Fig. 2c).

Together with the hierarchical orderings observed across sensory systems, these findings indicate that INTs systematically align with established structural markers of cortical hierarchy.

### Preserved cortical hierarchy of INTs in ASD

Building upon the preserved sensory hierarchy described above, we next examined whether the large-scale cortical hierarchy of INTs is similarly maintained across the whole brain in individuals with ASD. Vertex-, parcel-, and network-wise INTs were estimated for both TDC and individuals with ASD (Fig. 3). Group-averaged INT maps were computed from values adjusted for age, sex, and mean FD. The preserved spatial correspondence was clearly visible in the cortical surface maps, which displayed consistent gradients of INTs from primary sensorimotor to higher-order association regions in both ASD and TDC. Longer INTs were observed in association cortices, such as the default mode and frontoparietal networks, whereas shorter INTs appeared in primary sensory and motor regions. This hierarchical gradient mirrors previous findings in neurotypical populations (Wengler et al., 2020; Raut et al., 2020) and suggests that the global organization of temporal integration is maintained in ASD.

**Figure 3.**
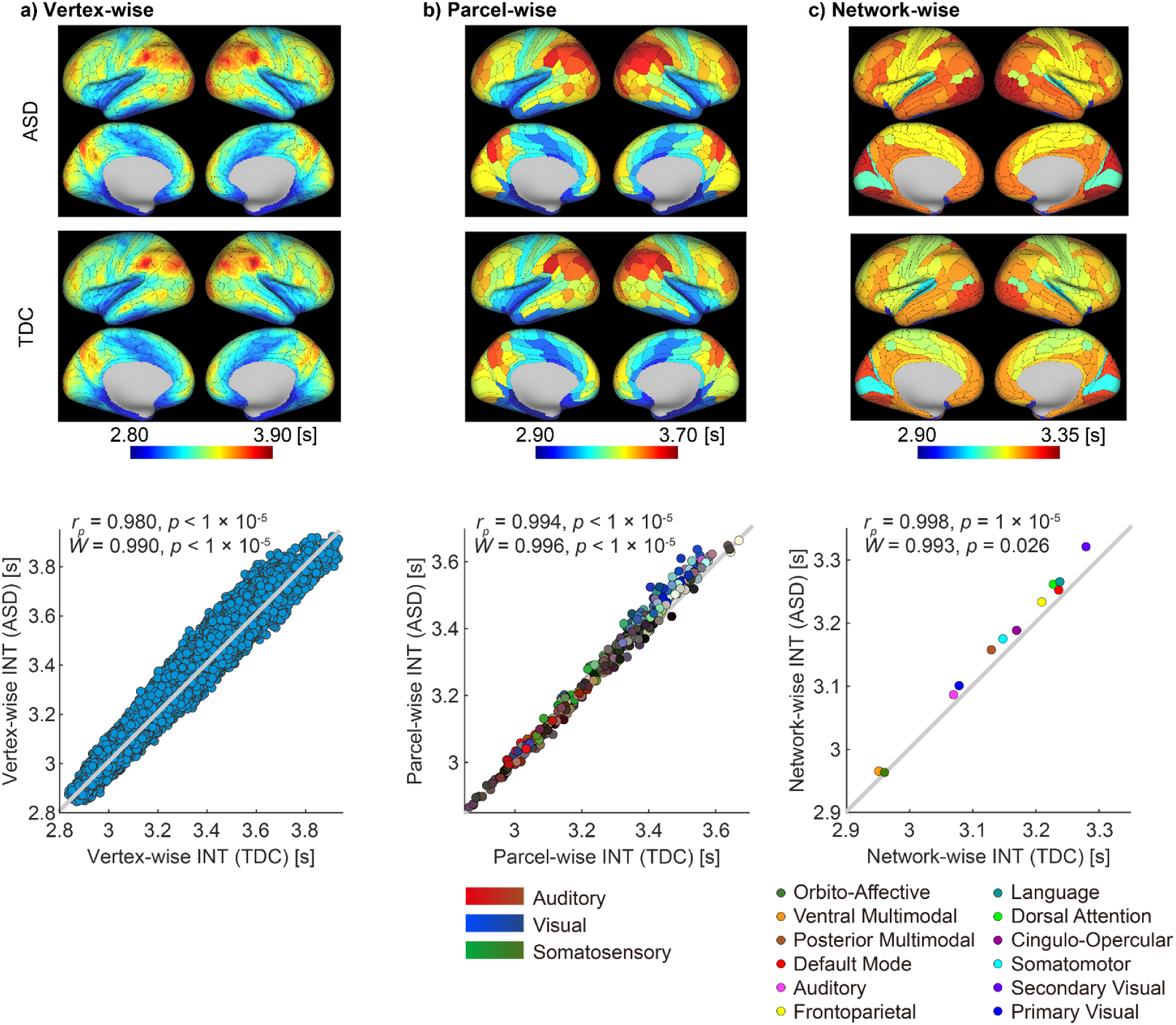
Preserved cortical hierarchy of intrinsic neural timescales (INTs) in autism spectrum disorder (ASD). Cortical surface maps show vertex-(a), parcel-(b), and network-wise (c) intrinsic neural timescales in ASD (top) and typically developed controls (TDC; middle), together with the corresponding cross-group comparisons (bottom). Scatter plots illustrate the correspondence of INTs between groups at each spatial scale. Each dot represents a vertex, cortical parcel, or functional network, and the gray diagonal line indicates perfect correspondence (Pearson’s correlation coefficient *r_p_* = 1) between ASD and TDC. Across all levels—from fine-grained vertices to large-scale functional networks—INTs exhibited an almost identical spatial distribution between ASD and TDC (vertex-wise: kendoll’s *W* = 0.990, *p* < 1×10⁻^5^; parcel-wise: *W* = 0.996, *p* < 1×10⁻^5^; network-wise: *W* = 0.993, *p* = 0.026). In panel (b), colors correspond to cortical regions defined in the Glasser multimodal parcellation (Glasser et al., 2016), whereas in panel (c), colors correspond to canonical large-scale functional networks based on the Cole–Anticevic brain-wide network partition (Ji et al., 2019).

To quantitatively assess this preservation, we compared the spatial correspondence of INTs between ASD and TDC across multiple scales. INTs exhibited remarkably similar spatial distributions between the two groups. Vertex-wise INTs showed extremely high concordance at the group level (Pearson’s correlation coefficient *r_p_* = 0.980, Kendall’s *W* = 0.990; both *p* < 1 × 10⁻^5^), indicating that both the absolute magnitude and the relative hierarchical order of cortical INTs were nearly identical between ASD and TDC. Comparable results were obtained at the parcel-wise scale (*r_p_*= 0.994, *W* = 0.996; both *p* < 1 × 10⁻^5^) and at the network-wise scale (*r_p_* = 0.998, p < 1 × 10⁻^5^; *W* = 0.993; *p* = 0.026), further confirming that the large-scale temporal hierarchy of the cortex is preserved in ASD.

Taken together, these results demonstrate that, beyond modality-specific hierarchies such as the auditory system, the overall cortical hierarchy of INTs is robustly preserved in ASD.

### Longer INTs are associated with larger group-related divergence

To further characterize how group differences vary across the preserved cortical hierarchy, we examined the relationship between the length of INTs and the magnitude of group differences (Cohen’s *d* for TDC–ASD) across multiple spatial scales (Fig. 4). All analyses were performed after adjusting for age, sex, and FD.

**Figure 4.**
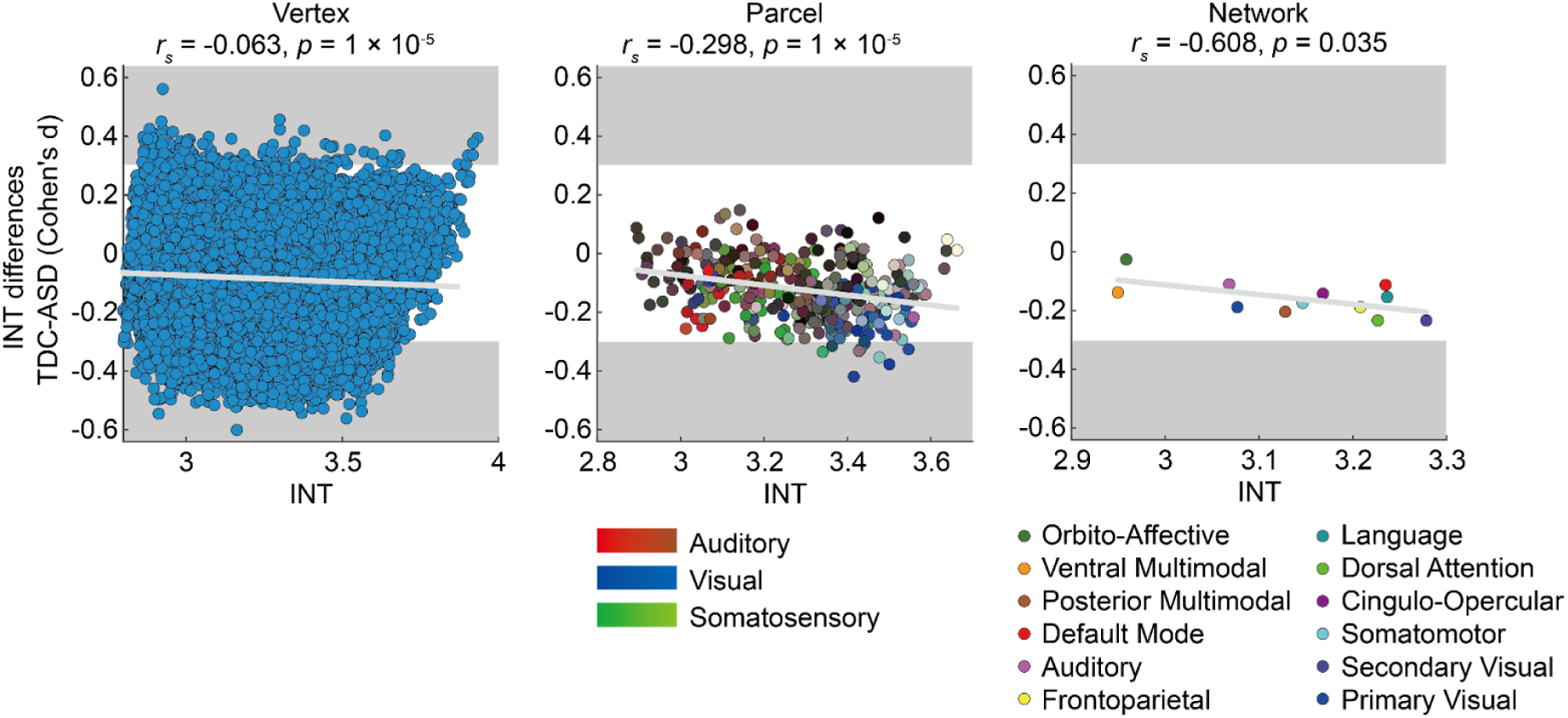
Longer intrinsic neural timescales (INTs) are associated with larger group differences between ASD and TDC. Scatter plots show the association between INTs in TDC (*x*-axis) and the corresponding group differences (Cohen’s *d* for TDC–ASD, *y*-axis) at the vertex (left), parcel (center), and network (right) levels. Each dot represents a cortical vertex, parcel, or network, respectively. The light gray line represents the Pearson’s correlation drawn for visualization, whereas the numerical values displayed at the top of each panel indicate the Spearman’s correlation coefficients (*r_s_*) quantifying the monotonic association between INTs and group differences. Colors in panel center correspond to cortical regions defined in the Glasser multimodal parcellation (Glasser et al., 2016), and colors in panel right correspond to canonical large-scale functional networks based on the Cole–Anticevic brain-wide network partition (Ji et al., 2019). Gray shading marks data points with uncorrected *p* < 0.05.

At all scales, no regions survived correction for multiple comparisons (FDR *p_FDR_* > 0.050) in the group-level analyses, indicating the absence of localized group effects. For transparency, parcels showing uncorrected effects at *p* < 0.05 are summarized in Table 2 and Supplementary Figure S1. Although no individual regions exhibited significant group differences after correction, a consistent trend was observed across the cortex: regions with longer INTs tended to show progressively larger TDC–ASD differences, reflecting prolongation of INTs in ASD along the global cortical timescale hierarchy. This negative association was evident at the vertex (Spearman’s *r_s_* = –0.063, *p* < 1 × 10^-5^), parcel (*r_s_* = –0.298, *p* < 1 × 10^-5^), and network (*r_s_* = –0.608, *p* = 0.036) levels.

**Table 2.** Parcels with significant between-group differences in intrinsic neural timescale identified by ANCOVA (typically developed control (TDC) versus autism spectrum disorder (ASD)), with age, sex and mean framewise displacement included as covariates. R, right; L, left; PIT, posterior inferior temporal area; LO2, lateral occipital area 2; MIP, medial intraparietal area; 7PC, part of Broadman area 7 in the superior parietal cortex; a9-46v, anterior 9-46 ventral; V3CD, visual area V3 complex dorsal subdivision; VIP, ventral intraparietal area; MT, middle temporal area; V4, visual area V4.

| Parcel name | Network label | $p$ value (unc.) | $p$ value (FDR) | ASD | TDC | Cohen's $d$ |
| --- | --- | --- | --- | --- | --- | --- |
| R_PIT | Visual2 | 0.007 | 0.822 | 3.531 | 3.417 | -0.420 |
| R_LO2 | Visual2 | 0.015 | 0.822 | 3.611 | 3.502 | -0.378 |
| L_MIP | Dorsal Attention | 0.022 | 0.822 | 3.560 | 3.478 | -0.355 |
| L_7PC | Somatomotor | 0.030 | 0.822 | 3.429 | 3.342 | -0.336 |
| L_a9-46v | Frontoparietal | 0.032 | 0.822 | 3.499 | 3.429 | -0.333 |
| R_V3CD | Visual2 | 0.035 | 0.822 | 3.637 | 3.548 | -0.327 |
| L_VIP | Visual2 | 0.036 | 0.822 | 3.516 | 3.434 | -0.324 |
| L_MT | Visual2 | 0.041 | 0.822 | 3.456 | 3.365 | -0.316 |
| R_V4 | Visual2 | 0.047 | 0.822 | 3.510 | 3.443 | -0.308 |

Together, these findings indicate that while no focal alterations were detected, the magnitude of group differences follows a global trend along the timescale hierarchy, pointing to a subtle modulation of long-timescale cortical processes in ASD.

### Individual deviations from the timescale hierarchy link sensory traits to cortical temporal dynamics

Finally, to examine whether individual variability in the cortical hierarchy of INTs is associated with behavioral traits, we analyzed the relationship between subject-specific deviations from the INT template and sensory processing profiles derived from the AASP questionnaire.

Because the AASP indexes subjective sensory experience rather than primary sensory thresholds, it reflects integrative and regulatory aspects of sensory processing that extend beyond early sensory encoding (Dunn, 1997, 2001). Such experiential dimensions may relate to variability in cortical temporal dynamics across hierarchical levels. Given that the large-scale hierarchical organization of INTs is preserved across individuals, inter-individual variability is expected to manifest primarily as deviations from this shared template rather than as alterations of the hierarchy itself. Quantifying subject-specific departures from the TDC-based INT template therefore provides a principled framework for characterizing individual variability while respecting the preserved hierarchical structure.

As shown in Fig. 5a, group-averaged INT profiles plotted against the parcel-wise mean INT of the TDC group, used as a template, exhibited a highly similar monotonic relationship in both ASD and TDC groups, indicating preservation of the overall hierarchical organization of INTs. To formally assess this observation, we fitted a subject-wise linear model aligning individual INT profiles to the template and compared the resulting parameters across groups. Neither the global offset parameter (α in eq.(1)), reflecting overall prolongation or shortening of INTs, nor the scaling parameter (*β*), reflecting the strength of hierarchical alignment to the template, differed significantly between ASD and TDC participants (both p>0.1). While these parameters showed no group-level differences, they were influenced by demographic and nuisance covariates, including age, sex, and head motion, underscoring the importance of accounting for such factors when characterizing individual INT profiles (Supplementary Table S1).

**Figure 5.**
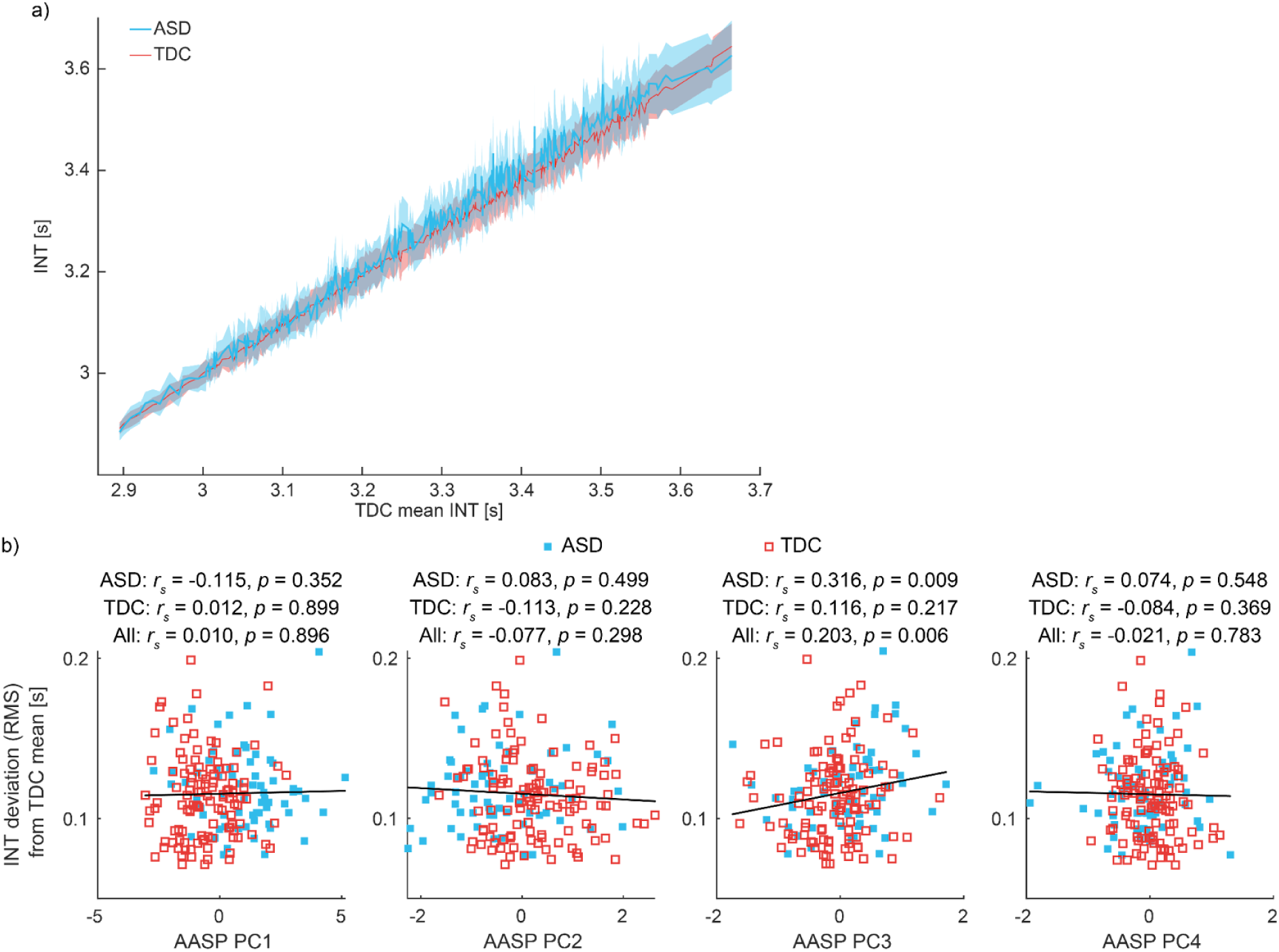
Individual deviations from the cortical hierarchy of intrinsic timescales and their relationship to sensory traits. (a) Group-averaged intrinsic neural timescale (INT) profiles plotted against the TDC-derived template across 360 cortical parcels. INT values were sorted according to the template ordering. Shaded areas represent 95% confidence intervals across participants. (b) Scatter plots showing relationships between individual deviation from the cortical timescale hierarchy, quantified as the root mean square (RMS) of residuals after linear alignment to the template, and principal component (PC) scores derived from the Adolescent/Adult Sensory Profile (AASP). PC1 reflects a general sensory reactivity dimension, PC2 is dominated by Sensation Seeking, PC3 contrasts Low Registration with Sensory Avoiding, and PC4 reflects opposing contributions of Sensory Sensitivity and Sensory Avoiding (see Table 3 for loadings).

**Table 3.** Group differences in Adolescent/Adult Sensory Profile (AASP) quadrant scores between individuals with autism spectrum disorder (ASD) and typically developed controls (TDC). Age and sex were included as covariates in the model to control for their potential confounding effects. Values in parentheses indicate the percentage of variance explained by each principal component (PC). Values in the PC columns represent component loadings.

|  | ASD Mean (SD) | TDC Mean (SD) | Group difference | PC1 (61.21 0%) | PC2 (24.02 2%) | PC3 (9.257 %) | PC4 (5.509 %) |
| --- | --- | --- | --- | --- | --- | --- | --- |
| Low registration | 39.388 (11.037) | 26.947 (8.009) | $t(178) = 8.553, p < 1 \times 10^{-5}$ | 0.552 | -0.084 | 0.798 | -0.226 |
| Sensation seeking | 34.298 (9.459) | 37.156 (9.025) | $t(178) = -1.644, p = 0.101$ | 0.182 | 0.976 | -0.053 | -0.107 |
| Sensory sensitivity | 40.925 (12.833) | 31.669 (9.056) | $t(178) = 5.535, p < 1 \times 10^{-5}$ | 0.589 | -0.034 | -0.189 | 0.785 |
| Sensation avoiding | 42.209 (10.608) | 32.191 (8.350) | $t(178) = 6.790, p < 1 \times 10^{-5}$ | 0.561 | -0.198 | -0.570 | -0.567 |

After accounting for global offset and hierarchical scaling, the residual RMS measure did not show a significant association with diagnostic groups or other demographic variables (Supplementary Table S1). Importantly, the absence of a group-level effect does not imply that this residual variability is unstructured; rather, it indicates that the variance captured by the residual RMS is not explained by diagnostic or demographic factors, making it well suited for probing continuous inter-individual differences that cut across diagnostic boundaries. To this end, we assessed relationships between the residual RMS measure and PC scores derived from the AASP, which were used to represent orthogonal sensory dimensions given the substantial correlations among AASP subscale raw scores, particularly among Low Registration, Sensory Sensitivity, and Sensation Avoiding (all *r_p_* > 0.60, p < 1 × 10^-5^). The PCA yielded four components with distinct loading patterns (Table 3): PC1 showed strong positive loadings on Low Registration, Sensory Sensitivity, and Sensation Avoiding, indicating a sensory modulation profile characterized by co-elevation across multiple response tendencies and elevated levels in ASD; PC2 was dominated by Sensation Seeking; PC3 exhibited a strong positive loading on Low Registration together with a substantial negative loading on Sensation Avoiding; PC4 was characterized by opposing contributions from Sensory Sensitivity and Sensory Avoiding. Correlation analyses were then performed between the residual RMS measure of INT and each AASP PC score.

As shown in Fig. 5b, partial correlation analyses controlling for age, sex, and FD revealed no significant associations between INT residual RMS and PC1, PC2, or PC4 in either group or in the combined sample (all *p* > 0.2). In contrast, a significant positive association emerged specifically for PC3. In the ASD group, higher PC3 scores were associated with larger residual RMS values (*r_s_* = 0.316, *p* = 0.009, *p_FDR_* = 0.037), and this relationship remained significant when all participants were considered together (*r_s_* = 0.203, *p* = 0.006, *p_FDR_* = 0.024). Although the association did not reach statistical significance in the TDC group alone, the direction of the effect was consistent across groups.

To assess the specificity of this effect, we conducted additional analysis using alternative INT metrics. First, we examined associations between raw INT values within the 12 functional networks and both AASP subscale scores and AASP PC scores. These exploratory analyses did not yield any correlations surviving FDR correction in any group (Supplementary Tables S2 and S3). Second, we tested whether a simpler deviation metric–defined as the direct difference between individual INT profiles and the template, without decomposing global offset and hierarchical scaling components–was associated with AASP PC scores. No significant associations were observed (all *p* > 0.080, *p_FDR_* > 0.250).

Taken together, these findings indicate that observed behavioral association is not explained by raw INT magnitude or network-specific values per se, but rather emerges specifically from individual deviations from the TDC-based INT template after accounting for global offset and hierarchical scaling.

## Discussion

In this study, we investigated whether the hierarchical organization of INT—a straightforward index characterizing the temporal profile of fMRI signals—is altered in ASD, and how individual variation along this hierarchy relates to sensory-processing traits. By quantifying INTs across four five-minute resting-state runs and situating each individual along the INT-based cortical hierarchy, we demonstrated that the overall hierarchical structure of INTs was preserved in ASD, while regions with longer INT exhibited further prolongation at the group level. Importantly, beyond global shifts and hierarchical scaling, we identified substantial individual-specific deviations from the canonical INT hierarchy. This pattern suggests that ASD is not characterized by a collapse of the hierarchical structure itself, but rather by systematic shifts and scaling biases that reflect demographic influences, alongside individual-specific deviations from this preserved template. These distributed deviations were selectively associated with sensory traits captured by the third principal component of the AASP, characterized by high Low Registration and low Sensory Avoiding, indicating that variability in sensory processing is linked not to absolute INT values but to individual departures from the INT-based cortical hierarchy of the cortex. Together, these findings indicate that while the macro-scale organization of cortical temporal dynamics remains largely intact in ASD, subtle individual-specific deviations—and their correspondence with sensory traits—reveal individual biases that may help explain heterogeneity in sensory experiences.

### Neurophysiological Implications of Prolonged Timescales

A prolonged intrinsic timescale can be interpreted as reflecting stronger local recurrent interactions and reduced influence from rapidly fluctuating inputs. Computational and empirical studies have shown that longer timescales emerge when local circuits exhibit stronger recurrent excitation, increased temporal integration, or a shift in the excitation–inhibition balance toward excitation. These mechanisms have been repeatedly implicated in ASD (Gogolla et al., 2009; Rubenstein & Merzenich, 2003; Trakoshis et al., 2020), where altered local connectivity, elevated cortical excitability, and atypical inhibitory signaling have been documented across multiple levels of analysis, including cellular, circuit, and macroscopic imaging studies (Gogolla et al., 2009; Rubenstein & Merzenich, 2003;

Takarae & Sweeney, 2017). Viewed in this context, the prolongation observed in longer-timescale cortical regions in our sample aligns well with these prior accounts. Our findings therefore suggest that, despite the preservation of the overall cortical temporal hierarchy, individuals with ASD may exhibit subtle but systematic shifts in local circuit dynamics that bias certain regions toward slower intrinsic activity patterns. Such regional prolongation may reflect underlying circuit-level alterations that contribute to the sensory and cognitive characteristics associated with the condition.

### Discrepancy with reports indicating shortened INT in ASD

Several studies have reported locally shortened INTs in ASD (Uscătescu et al., 2023; Watanabe et al., 2019), whereas our findings indicate a largely preserved hierarchical organization of INTs at the group level. A key factor that may contribute to these discrepancies is the substantial heterogeneity within ASD, which increases the likelihood that individual studies—each sampling a different part of the spectrum—capture distinct patterns of cortical dynamics (Lenroot & Yeung, 2013; Pelphrey et al., 2011; Tang et al., 2020). This issue is further amplified by sample-size–dependent variability: small samples are known to produce unstable estimates, including effects that can occasionally reverse in sign relative to the population-level pattern (Marek et al., 2022). To enhance the stability of intrinsic timescale estimation, we designed the acquisition and preprocessing pipeline to minimize motion-related confounds and sampling variability, including multiple resting-state runs and conservative motion control criteria. This approach may have contributed to the stability of the observed hierarchical organization.

Despite these efforts to improve estimation stability, larger samples would, in principle, further mitigate sampling fluctuations. However, expanding sample size in clinical neuroimaging is challenging. Data quality can be compromised by increased head motion, some individuals may decline or discontinue scanning due to sensory sensitivities, scanner noise, physical restraint, or anxiety, and comorbid psychiatric conditions (e.g., depression or panic symptoms) often limit eligibility for MRI participation. These considerations underscore the need for large, harmonized datasets and, ultimately, meta-analytic integration to determine which INT alterations are robust and which are cohort-specific.

Beyond sampling-related factors, it is also important to consider how INT should be interpreted as a signal-derived metric. Although INT is often discussed as reflecting the temporal window of neural information integration, it is formally estimated from measured neural signals and therefore depends not only on underlying neural dynamics but also on signal quality. Even if a neural system operates with relatively long intrinsic integration windows, increased noise, reduced signal-to-noise ratio, or motion-related artifacts can bias the estimated autocorrelation structure toward shorter timescales. Consequently, a shortened INT does not necessarily imply a genuine reduction in the temporal window of information processing.

In this sense, INT may be better understood as an index constrained by structural and physiological properties of cortical circuits, rather than a direct readout of functional integration timescales. While this makes INT valuable for characterizing large-scale hierarchical organization and inter-regional differences, caution is warranted when interpreting individual-level correlations or attributing changes in INT solely to alterations in cognitive or perceptual processing.

### Sensory traits relate to individual deviations from the cortical INT hierarchy

Beyond group-averaged effects, a central contribution of the present study lies in the characterization of individual-specific deviations from the INT-based cortical hierarchy. By explicitly modeling and removing global shifts and hierarchical scaling, we isolated distributed residual variability that captures how each individual departs from the canonical temporal organization of the cortex. This residual-based measure revealed meaningful associations with sensory traits, despite the absence of robust correlations between raw INT values and sensory measures at the level of individual networks. Together, these findings indicate that associations with sensory traits are not driven by absolute intrinsic timescale values at specific regions, but instead relate to distributed deviations from the INT-based cortical hierarchy. This observation is consistent with the view that sensory experience reflects coordinated processing across cortical regions rather than isolated regional dynamics (Pessoa, 2014, 2018; Uddin et al., 2013).

The association between residual INT deviation and the AASP principal component contrasting higher Low Registration with lower Sensory Avoiding underscores the relevance of a distributed perspective on sensory traits. This component captures a sensory profile characterized by reduced sensory registration accompanied by lower tendencies toward avoidance, independent of the shared variance among AASP subscales (largely reflected in PC1). The selective relationship with residual INT deviation suggests that such sensory profiles are linked not to absolute intrinsic timescale values in specific regions, but to how temporal processing is unevenly organized across the cortical hierarchy. In this context, increased deviation from the INT-based cortical hierarchy may reflect an altered balance between local temporal integration and hierarchical coordination, giving rise to idiosyncratic sensory experiences that are not well captured by group-level averages.

Importantly, the fact that this component reflects reduced sensory registration rather than heightened sensory sensitivity provides a plausible mechanistic link to temporal processing. Reduced sensory registration may require sensory evidence to be accumulated over longer temporal windows before reliably influencing perceptual experience (Gold & Shadlen, 2007; Wang, 2002), thereby rendering such profiles particularly sensitive to how temporal integration is distributed across cortical levels. By contrast, sensory hypersensitivity may arise from increased background activity or reduced signal-to-noise ratio in early sensory regions—mechanisms that may influence perceptual responsiveness without necessarily altering the large-scale distribution of temporal integration across cortical levels (Gerstner et al., 2014). Taken together, these considerations support the interpretation that residual deviations from the INT-based cortical hierarchy are consistent with alterations in temporal accumulation across processing levels, rather than general sensory responsiveness.

It is also worth noting that, while global offset and hierarchical scaling parameters did not differ between diagnostic groups, they showed systematic associations with demographic factors. In particular, female participants tended to exhibit overall longer INTs and a shallower hierarchical slope. However, given the relatively small number of female participants in the present sample, these effects should be interpreted cautiously and warrant further investigation in larger, sex-balanced cohorts.

More broadly, these results align with emerging views that emphasize ASD as a condition characterized not by a single canonical neural alteration, but by pronounced inter-individual variability in neural organization. From this perspective, deviations from normative hierarchical patterns—rather than shifts in the hierarchy itself—may constitute a key neural substrate of phenotypic heterogeneity. Residual-based metrics, such as those introduced here, offer a complementary approach to traditional group comparisons by directly quantifying individual differences in large-scale cortical dynamics, and may prove useful for linking intrinsic brain organization to behavioral diversity in both clinical and non-clinical populations.

### Limitation

Although no statistically significant difference was observed between the ASD and TDC groups in the auditory INT gradient, the direction of the effect was consistent with previous findings suggesting slightly reduced temporal integration in ASD (Uscătescu et al., 2023; Watanabe et al., 2019). Based on the observed group effect in the present data (Cohen’s *d* ≈ 0.21), detecting such a difference with 80% (90%) power would require approximately 233 (311) participants per group. More generally, increasing sample sizes remains an important consideration in neuroimaging studies to reliably characterize subtle group-level effects. At the same time, the small magnitude of the estimated group difference suggests that any potential alteration in auditory temporal hierarchy, if present, is likely modest.

A further limitation concerns the substantial heterogeneity within the ASD group (Lenroot & Yeung, 2013; Masi et al., 2017; Uljarević et al., 2017). Sensory-processing traits showed wide inter-individual variability, raising the possibility that subgroup-specific characteristics may influence the overall group estimates of INT, particularly in slow-timescale regions.

Future studies with larger samples or stratification approaches may help clarify whether distinct sensory phenotypes exhibit different hierarchical signatures of INT.

Taken together, these considerations highlight the need for future work that integrates larger samples and multimodal neural markers to more precisely characterize how cortical temporal hierarchies vary across individuals and contribute to sensory experiences.

## Author Contributions

**Yumi Shikauch**: Conceptualization, Software, Formal analysis, Writing - Original Draft, Visualization. **Ryuta Aoki**: Conceptualization, Methodology, Investigation, Data Curation, Writing - Review & Editing, Supervision. **Takashi Itahashi**: Methodology, Software, Investigation, Data Curation, Writing - Review & Editing. **Masaaki Shimizu**: Investigation, Data Curation, Writing - Review & Editing. **Taiga Naoe**: Writing - Review & Editing. **Tsukasa Okimura**: Resources, Writing - Review & Editing. **Haruhisa Ohta**: Resources, Writing - Review & Editing. **Ryu-ichiro Hashimoto**: Methodology, Investigation, Writing - Review & Editing, Funding acquisition. **Motoaki Nakamura**: Conceptualization, Investigation, Resources, Writing - Review & Editing, Supervision, Project administration, Funding acquisition.

## Acknowledgements

We would like to express our gratitude to Noriko Ishimura, Mika Kato, and Taku Sato for their contributions to recruitment of participants for this study.

## Supplementary Materials

**Supplementary Figure S1.**
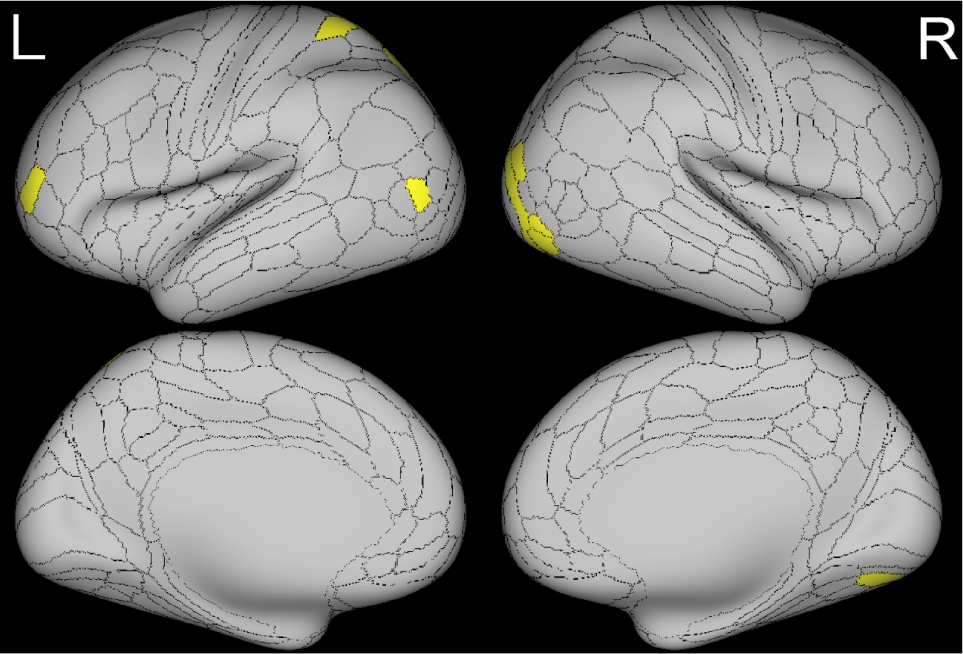
Spatial distribution of parcels showing group-related divergence in intrinsic neural timescales. Binary cortical surface maps illustrating the anatomical locations of Glasser parcels that exhibited group-related divergence between the ASD and TDC groups. Parcels showing uncorrected significance (*p* < 0.05) in the parcel-wise comparison are highlighted in yellow, while all other parcels are shown in gray.

---

**Supplementary Table S1.** Effects of diagnostic group and covariates on individual deviation metrics derived from intrinsic neural timescale (INT) profiles. ANCOVA models included group (ASD vs. TDC), age, sex, and mean framewise displacement (FD) as predictors. The global offset parameter (α) reflects overall prolongation or shortening of INTs (a), the scaling parameter (*β*) reflects hierarchical alignment to the template (b), and rms quantifies individual-specific deviations from the INT-based cortical hierarchy after accounting for global and hierarchical effects (c). Significant effects (*p* < 0.05) are highlighted in yellow.

| Predictor | Estimate | SE | $t$ | $p$ |
| --- | --- | --- | --- | --- |
| Group (TDC) | 0.176 | 0.120 | 1.460 | 0.145 |
| Age | -0.017 | 0.008 | -2.070 | 0.040 |
| Sex (Female) | 0.623 | 0.130 | 4.810 | $< 1 \times 10^{-5}$ |
| FD | 4.957 | 2.281 | 2.170 | 0.031 |

b) Hierarchical scaling parameter ( $\beta$ ) Model statistics: $F(4,177) = 7.329$ , $p = 1.744 \times 10^{-5}$
| Predictor | Estimate | SE | $t$ | $p$ |
| --- | --- | --- | --- | --- |
| Group (TDC) | -0.062 | 0.042 | -1.456 | 0.147 |
| Age | 0.004 | 0.003 | 1.436 | 0.153 |
| Sex (Female) | -0.212 | 0.046 | -4.634 | $< 1 \times 10^{-5}$ |
| FD | -1.499 | 0.805 | -1.864 | 0.064 |

c) Residual deviation from template ( $r_{ms}$ )891 Model statistics: $F(4,177) = 0.828$ , $p = 0.508$
| Predictor | Estimate | SE | $t$ | $p$ |
| --- | --- | --- | --- | --- |
| Group (TDC) | -0.006 | 0.004 | -1.570 | 0.119 |
| Age | 0.000 | 0.000 | 0.400 | 0.690 |
| Sex (Female) | -0.003 | 0.004 | -0.750 | 0.453 |
| FD | -0.048 | 0.077 | -0.620 | 0.535 |

**Supplementary Table S2.** Spearman correlations between intrinsic neural timescales (INT) and Adolescent/Adult Sensory Profile (AASP) subscale scores across functional networks. Spearman’s rank correlation coefficients (*r_s_*), uncorrected p-values (*p_unc_*) are reported for each AASP subscale (Low Registration, Sensation Seeking, Sensory Sensitivity, Sensation Avoiding) within each functional network separately for ASD, TDC, and the combined participant (All). Cells highlighted in yellow indicate associations with uncorrected *p* < 0.05. No correlations survived FDR correction (all > 0.700).

|  |  | Low Registration |  | Sensation Seeking |  | Sensory Sensitivity |  | Sensation Avoiding |  |
| --- | --- | --- | --- | --- | --- | --- | --- | --- | --- |
| | | $r_s$ | $p$ | $r_s$ | $p$ | $r_s$ | $p$ | $r_s$ | $p$ |
| Visual | ASD | 0.083 | 0.504 | 0.073 | 0.555 | -0.025 | 0.842 | -0.097 | 0.437 |
|  | TDC | 0.047 | 0.620 | -0.166 | 0.077 | 0.028 | 0.770 | 0.062 | 0.507 |
|  | All | 0.097 | 0.194 | -0.073 | 0.328 | 0.019 | 0.794 | 0.025 | 0.733 |
| Visual2 | ASD | 0.000 | 0.997 | 0.089 | 0.476 | -0.091 | 0.462 | -0.136 | 0.272 |
|  | TDC | 0.065 | 0.492 | -0.174 | 0.063 | 0.013 | 0.889 | 0.030 | 0.749 |
|  | All | 0.084 | 0.257 | -0.076 | 0.311 | -0.004 | 0.961 | 0.004 | 0.958 |
| Somatomotor | ASD | 0.018 | 0.883 | 0.102 | 0.410 | -0.127 | 0.306 | -0.208 | 0.092 |
|  | TDC | 0.027 | 0.779 | -0.116 | 0.216 | 0.000 | 0.997 | -0.054 | 0.568 |
|  | All | 0.061 | 0.414 | -0.031 | 0.676 | -0.024 | 0.752 | -0.080 | 0.282 |
| Cingulo-Opercular | ASD | 0.076 | 0.539 | 0.054 | 0.664 | -0.090 | 0.469 | -0.122 | 0.327 |
|  | TDC | 0.044 | 0.642 | -0.122 | 0.194 | 0.057 | 0.545 | -0.002 | 0.979 |
|  | All | 0.067 | 0.371 | -0.052 | 0.482 | 0.006 | 0.937 | -0.038 | 0.614 |
| Dorsal Attention | ASD | 0.044 | 0.725 | 0.146 | 0.239 | -0.092 | 0.460 | -0.122 | 0.325 |
|  | TDC | 0.022 | 0.815 | -0.158 | 0.092 | 0.037 | 0.697 | -0.009 | 0.920 |
|  | All | 0.065 | 0.383 | -0.048 | 0.522 | 0.012 | 0.876 | -0.018 | 0.812 |
| Language | ASD | 0.046 | 0.712 | 0.045 | 0.720 | -0.117 | 0.344 | -0.157 | 0.204 |
|  | TDC | 0.056 | 0.551 | -0.137 | 0.144 | 0.093 | 0.322 | 0.079 | 0.399 |
|  | All | 0.074 | 0.323 | -0.062 | 0.405 | 0.025 | 0.738 | 0.008 | 0.920 |
| Frontoparietal | ASD | 0.079 | 0.527 | 0.076 | 0.543 | -0.104 | 0.401 | -0.088 | 0.479 |
|  | TDC | 0.021 | 0.827 | -0.154 | 0.099 | 0.052 | 0.583 | 0.049 | 0.605 |
|  | All | 0.062 | 0.407 | -0.063 | 0.395 | 0.003 | 0.973 | 0.010 | 0.894 |
| Auditory | ASD | -0.002 | 0.988 | 0.054 | 0.663 | -0.154 | 0.213 | -0.251 | 0.041 |
|  | TDC | 0.044 | 0.639 | -0.160 | 0.088 | 0.043 | 0.647 | -0.023 | 0.806 |
|  | All | 0.054 | 0.470 | -0.069 | 0.358 | -0.024 | 0.748 | -0.081 | 0.279 |
| Default | ASD | 0.067 | 0.588 | 0.094 | 0.451 | -0.110 | 0.375 | -0.071 | 0.568 |
|  | TDC | 0.053 | 0.573 | -0.122 | 0.193 | 0.073 | 0.438 | 0.041 | 0.665 |
|  | All | 0.061 | 0.411 | -0.028 | 0.705 | 0.004 | 0.961 | -0.002 | 0.981 |
| Posterior Multimodal | ASD | 0.016 | 0.896 | 0.060 | 0.629 | -0.110 | 0.376 | -0.184 | 0.135 |
|  | TDC | 0.047 | 0.620 | -0.150 | 0.109 | 0.040 | 0.669 | -0.001 | 0.989 |
|  | All | 0.075 | 0.317 | -0.072 | 0.337 | -0.001 | 0.988 | -0.038 | 0.609 |
| Ventral Multimodal | ASD | 0.087 | 0.483 | 0.048 | 0.697 | -0.001 | 0.996 | -0.114 | 0.360 |
|  | TDC | 0.082 | 0.384 | -0.139 | 0.138 | 0.055 | 0.563 | 0.037 | 0.693 |
|  | All | 0.113 | 0.129 | -0.069 | 0.356 | 0.045 | 0.545 | -0.002 | 0.977 |
| Orbito-Affective | ASD | 0.024 | 0.847 | 0.045 | 0.720 | -0.056 | 0.655 | -0.127 | 0.307 |
|  | TDC | 0.110 | 0.244 | -0.140 | 0.136 | 0.070 | 0.460 | 0.026 | 0.782 |
|  | All | 0.059 | 0.428 | -0.055 | 0.459 | -0.006 | 0.932 | -0.061 | 0.415 |

**Supplementary Table S3.** Spearman correlations between intrinsic neural timescales (INT) and principal component (PC) scores derived from the Adolescent/Adult Sensory Profile (AASP) across functional networks. Spearman’s rank correlation coefficients (*r_s_*) and uncorrected *p*-values are shown for each AASP principal component across functional networks, analyzed separately in ASD, TDC, and the combined participant sample (All). Principal component analysis was applied to the four AASP subscales to obtain orthogonal sensory dimensions capturing shared and distinct variance across subscales. Cells highlighted in yellow indicate associations with uncorrected p < 0.05. No associations remained significant after false discovery rate (FDR) correction (all > 0.08).

|  |  | PC1 |  | PC2 |  | PC3 |  | PC4 |  |
| --- | --- | --- | --- | --- | --- | --- | --- | --- | --- |
| | | $r_s$ | $p$ | $r_s$ | $p$ | $r_s$ | $p$ | $r_s$ | $p$ |
| Visual | ASD | -0.003 | 0.981 | 0.073 | 0.558 | 0.315 | 0.010 | 0.092 | 0.458 |
|  | TDC | 0.015 | 0.874 | -0.197 | 0.035 | 0.038 | 0.684 | 0.022 | 0.815 |
|  | All | 0.035 | 0.638 | -0.103 | 0.168 | 0.160 | 0.032 | 0.029 | 0.696 |
| Visual2 | ASD | -0.070 | 0.571 | 0.116 | 0.350 | 0.259 | 0.034 | 0.056 | 0.652 |
|  | TDC | 0.009 | 0.927 | -0.199 | 0.033 | 0.097 | 0.304 | 0.007 | 0.941 |
|  | All | 0.021 | 0.781 | -0.095 | 0.201 | 0.166 | 0.025 | 0.015 | 0.842 |
| Somatomotor | ASD | -0.109 | 0.379 | 0.149 | 0.229 | 0.337 | 0.006 | 0.080 | 0.519 |
|  | TDC | -0.027 | 0.770 | -0.121 | 0.199 | 0.114 | 0.224 | 0.108 | 0.251 |
|  | All | -0.019 | 0.801 | -0.021 | 0.781 | 0.201 | 0.007 | 0.092 | 0.217 |
| Cingulo-Opercular | ASD | -0.059 | 0.636 | 0.061 | 0.622 | 0.296 | 0.015 | 0.035 | 0.779 |
|  | TDC | 0.021 | 0.827 | -0.141 | 0.133 | 0.097 | 0.304 | 0.128 | 0.173 |
|  | All | 0.009 | 0.906 | -0.057 | 0.442 | 0.159 | 0.032 | 0.095 | 0.202 |
| Dorsal Attention | ASD | -0.045 | 0.715 | 0.161 | 0.193 | 0.294 | 0.016 | 0.038 | 0.760 |
|  | TDC | -0.002 | 0.983 | -0.174 | 0.063 | 0.087 | 0.355 | 0.106 | 0.258 |
|  | All | 0.020 | 0.790 | -0.057 | 0.441 | 0.158 | 0.033 | 0.078 | 0.297 |
| Language | ASD | -0.077 | 0.537 | 0.060 | 0.628 | 0.323 | 0.008 | 0.048 | 0.698 |
|  | TDC | 0.071 | 0.453 | -0.168 | 0.073 | 0.025 | 0.790 | 0.081 | 0.391 |
|  | All | 0.037 | 0.622 | -0.075 | 0.314 | 0.135 | 0.069 | 0.067 | 0.368 |
| Frontoparietal | ASD | -0.033 | 0.790 | 0.081 | 0.512 | 0.282 | 0.021 | -0.009 | 0.942 |
|  | TDC | 0.023 | 0.810 | -0.182 | 0.052 | 0.032 | 0.737 | 0.059 | 0.530 |
|  | All | 0.021 | 0.780 | -0.078 | 0.295 | 0.130 | 0.081 | 0.032 | 0.664 |
| Auditory | ASD | -0.144 | 0.245 | 0.122 | 0.323 | 0.355 | 0.003 | 0.045 | 0.719 |
|  | TDC | -0.004 | 0.967 | -0.173 | 0.065 | 0.106 | 0.259 | 0.144 | 0.124 |
|  | All | -0.029 | 0.695 | -0.054 | 0.469 | 0.192 | 0.010 | 0.096 | 0.197 |
| Default | ASD | -0.031 | 0.800 | 0.089 | 0.474 | 0.264 | 0.031 | -0.056 | 0.653 |
|  | TDC | 0.039 | 0.677 | -0.144 | 0.125 | 0.053 | 0.574 | 0.072 | 0.442 |
|  | All | 0.017 | 0.825 | -0.039 | 0.600 | 0.129 | 0.083 | 0.027 | 0.718 |
| Posterior Multimodal | ASD | -0.097 | 0.435 | 0.099 | 0.423 | 0.338 | 0.005 | 0.050 | 0.688 |
|  | TDC | 0.019 | 0.841 | -0.170 | 0.069 | 0.101 | 0.282 | 0.088 | 0.351 |
|  | All | 0.008 | 0.914 | -0.073 | 0.329 | 0.186 | 0.012 | 0.068 | 0.364 |
| Ventral Multimodal | ASD | 0.006 | 0.959 | 0.069 | 0.576 | 0.306 | 0.012 | 0.140 | 0.257 |
|  | TDC | 0.041 | 0.661 | -0.169 | 0.070 | 0.081 | 0.387 | 0.067 | 0.479 |
|  | All | 0.048 | 0.517 | -0.082 | 0.271 | 0.169 | 0.023 | 0.089 | 0.233 |
| Orbito-Affective | ASD | -0.045 | 0.715 | 0.070 | 0.575 | 0.256 | 0.037 | 0.106 | 0.395 |
|  | TDC | 0.044 | 0.642 | -0.161 | 0.086 | 0.114 | 0.225 | 0.100 | 0.286 |
|  | All | -0.015 | 0.840 | -0.047 | 0.525 | 0.164 | 0.027 | 0.121 | 0.105 |

## References

1. Behzadi, Y., Restom, K., Liau, J., & Liu, T. T. (2007). A component based noise correction method (CompCor) for BOLD and perfusion based fMRI. NeuroImage, 37(1), 90–101.

2. Bernhardt, B. C., Valk, S. L., Hong, S.-J., Soulières, I., & Mottron, L. (2025). Autism-related shifts in the brain’s information processing hierarchy. Trends in Cognitive Sciences, 29(10), 942–955.

3. Brown, C., Tollefson, N., Dunn, W., Cromwell, R., & Filion, D. (2001). The Adult Sensory Profile: measuring patterns of sensory processing. The American Journal of Occupational Therapy: Official Publication of the American Occupational Therapy Association, 55(1), 75–82.

4. Burt, J. B., Demirtaş, M., Eckner, W. J., Navejar, N. M., Ji, J. L., Martin, W. J., Bernacchia, A., Anticevic, A., & Murray, J. D. (2018). Hierarchy of transcriptomic specialization across human cortex captured by structural neuroimaging topography. Nature Neuroscience, 21(9), 1251–1259.

5. Chaudhuri, R., Knoblauch, K., Gariel, M.-A., Kennedy, H., & Wang, X.-J. (2015). A large-scale circuit mechanism for hierarchical dynamical processing in the primate cortex. Neuron, 88(2), 419–431.

6. Dunn, W. (1997). The impact of sensory processing abilities on the daily lives of young children and their families: A conceptual model. Infants and Young Children, 9(4), 23–35.

7. Dunn, W. (2001). The sensations of everyday life: empirical, theoretical, and pragmatic considerations. The American Journal of Occupational Therapy: Official Publication of the American Occupational Therapy Association, 55(6), 608–620.

8. Fukutomi, H., Glasser, M. F., Zhang, H., Autio, J. A., Coalson, T. S., Okada, T., Togashi, K., Van Essen, D. C., & Hayashi, T. (2018). Neurite imaging reveals microstructural variations in human cerebral cortical gray matter. NeuroImage, 182, 488–499.

9. Gerstner, W., Kistler, W. M., Naud, R., & Paninski, L. (2014). Neuronal dynamics: From single neurons to networks and models of cognition. Cambridge University Press.

10. Glasser, M. F., Coalson, T. S., Robinson, E. C., Hacker, C. D., Harwell, J., Yacoub, E., Ugurbil, K., Andersson, J., Beckmann, C. F., Jenkinson, M., Smith, S. M., & Van Essen, D. C. (2016). A multi-modal parcellation of human cerebral cortex. Nature, 536(7615), 171–178.

11. Glasser, M. F., Sotiropoulos, S. N., Wilson, J. A., Coalson, T. S., Fischl, B., Andersson, J. L., Xu, J., Jbabdi, S., Webster, M., Polimeni, J. R., Van Essen, D. C., Jenkinson, M., & WU-Minn HCP Consortium. (2013). The minimal preprocessing pipelines for the Human Connectome Project. NeuroImage, 80, 105–124.

12. Glasser, M. F., & Van Essen, D. C. (2011). Mapping human cortical areas in vivo based on myelin content as revealed by T1- and T2-weighted MRI. The Journal of Neuroscience: The Official Journal of the Society for Neuroscience, 31(32), 11597–11616.

13. Gogolla, N., Leblanc, J. J., Quast, K. B., Südhof, T. C., Fagiolini, M., & Hensch, T. K. (2009). Common circuit defect of excitatory-inhibitory balance in mouse models of autism. Journal of Neurodevelopmental Disorders, 1(2), 172–181.

14. Gold, J. I., & Shadlen, M. N. (2007). The neural basis of decision making. Annual Review of Neuroscience, 30(1), 535–574.

15. Golesorkhi, M., Gomez-Pilar, J., Zilio, F., Berberian, N., Wolff, A., Yagoub, M. C. E., & Northoff, G. (2021). The brain and its time: intrinsic neural timescales are key for input processing. Communications Biology, 4(1), 970.

16. Honey, C. J., Thesen, T., Donner, T. H., Silbert, L. J., Carlson, C. E., Devinsky, O., Doyle, W. K., Rubin, N., Heeger, D. J., & Hasson, U. (2012). Slow cortical dynamics and the accumulation of information over long timescales. Neuron, 76(2), 423–434.

17. Huntenburg, J. M., Bazin, P.-L., & Margulies, D. S. (2018). Large-scale gradients in human cortical organization. Trends in Cognitive Sciences, 22(1), 21–31.

18. Itahashi, T., Tanji, K., Shikauchi, Y., Naoe, T., Okimura, T., Nakamura, M., Ohta, H., & Hashimoto, R.-I. (2025). Iron deposition and functional connectivity alterations in the right substantia nigra of adult males with autism. Cerebral Cortex (New York, N.Y.: 1991), 35(8), bhaf216.

19. Ito, T., Hearne, L. J., & Cole, M. W. (2020). A cortical hierarchy of localized and distributed processes revealed via dissociation of task activations, connectivity changes, and intrinsic timescales. NeuroImage, 221(117141), 117141.

20. Ji, J. L., Spronk, M., Kulkarni, K., Repovš, G., Anticevic, A., & Cole, M. W. (2019). Mapping the human brain’s cortical-subcortical functional network organization. NeuroImage, 185, 35–57.

21. Koike, S., Tanaka, S. C., Okada, T., Aso, T., Yamashita, A., Yamashita, O., Asano, M., Maikusa, N., Morita, K., Okada, N., Fukunaga, M., Uematsu, A., Togo, H., Miyazaki, A., Murata, K., Urushibata, Y., Autio, J., Ose, T., Yoshimoto, J.,…Hayashi, T. (2021). Brain/MINDS beyond human brain MRI project: A protocol for multi-level harmonization across brain disorders throughout the lifespan. NeuroImage: Clinical, 30, 102600.

22. Lenroot, R. K., & Yeung, P. K. (2013). Heterogeneity within autism spectrum disorders: What have we learned from neuroimaging studies? Frontiers in Human Neuroscience, 7, 733.

23. Marek, S., Tervo-Clemmens, B., Calabro, F. J., Montez, D. F., Kay, B. P., Hatoum, A. S., Donohue, M. R., Foran, W., Miller, R. L., Hendrickson, T. J., Malone, S. M., Kandala, S., Feczko, E., Miranda-Dominguez, O., Graham, A. M., Earl, E. A., Perrone, A. J., Cordova, M., Doyle, O.,…Dosenbach, N. U. F. (2022). Reproducible brain-wide association studies require thousands of individuals. Nature, 603(7902), 654–660.

24. Masi, A., DeMayo, M. M., Glozier, N., & Guastella, A. J. (2017). An overview of autism spectrum disorder, heterogeneity and treatment options. Neuroscience Bulletin, 33(2), 183–193.

25. Murray, J. D., Bernacchia, A., Freedman, D. J., Romo, R., Wallis, J. D., Cai, X., Padoa-Schioppa, C., Pasternak, T., Seo, H., Lee, D., & Wang, X.-J. (2014). A hierarchy of intrinsic timescales across primate cortex. Nature Neuroscience, 17(12), 1661–1663.

26. Paquola, C., Vos De Wael, R., Wagstyl, K., Bethlehem, R. A. I., Hong, S.-J., Seidlitz, J., Bullmore, E. T., Evans, A. C., Misic, B., Margulies, D. S., Smallwood, J., & Bernhardt, B. C. (2019). Microstructural and functional gradients are increasingly dissociated in transmodal cortices. PLoS Biology, 17(5), e3000284.

27. Pelphrey, K. A., Shultz, S., Hudac, C. M., & Vander Wyk, B. C. (2011). Research review: Constraining heterogeneity: the social brain and its development in autism spectrum disorder: Research Review: Constraining heterogeneity. Journal of Child Psychology and Psychiatry, and Allied Disciplines, 52(6), 631–644.

28. Pessoa, L. (2014). Understanding brain networks and brain organization. Physics of Life Reviews, 11(3), 400–435.

29. Pessoa, L. (2018). Understanding emotion with brain networks. Current Opinion in Behavioral Sciences, 19, 19–25.

30. Power, J. D., Barnes, K. A., Snyder, A. Z., Schlaggar, B. L., & Petersen, S. E. (2012). Spurious but systematic correlations in functional connectivity MRI networks arise from subject motion. NeuroImage, 59(3), 2142–2154.

31. Raut, R. V., Snyder, A. Z., & Raichle, M. E. (2020). Hierarchical dynamics as a macroscopic organizing principle of the human brain. Proceedings of the National Academy of Sciences of the United States of America, 117(34), 20890–20897.

32. Rubenstein, J. L. R., & Merzenich, M. M. (2003). Model of autism: increased ratio of excitation/inhibition in key neural systems: Model of autism. Genes, Brain, and Behavior, 2(5), 255–267.

33. Runyan, C. A., Piasini, E., Panzeri, S., & Harvey, C. D. (2017). Distinct timescales of population coding across cortex. Nature, 548(7665), 92–96.

34. Takarae, Y., & Sweeney, J. (2017). Neural hyperexcitability in autism spectrum disorders. Brain Sciences, 7(10), 129.

35. Tang, S., Sun, N., Floris, D. L., Zhang, X., Di Martino, A., & Yeo, B. T. T. (2020). Reconciling dimensional and categorical models of autism heterogeneity: A brain connectomics and behavioral study. Biological Psychiatry, 87(12), 1071–1082.

36. Trakoshis, S., Martínez-Cañada, P., Rocchi, F., Canella, C., You, W., Chakrabarti, B., Ruigrok, A. N., Bullmore, E. T., Suckling, J., Markicevic, M., Zerbi, V., MRC AIMS Consortium, Baron-Cohen, S., Gozzi, A., Lai, M.-C., Panzeri, S., & Lombardo, M. V. (2020). Intrinsic excitation-inhibition imbalance affects medial prefrontal cortex differently in autistic men versus women. eLife, 9. 10.7554/eLife.55684

37. Turjeman-Levi, Y., & Kluger, A. N. (2022). Sensory-processing sensitivity versus the sensory-processing theory: Convergence and divergence. Frontiers in Psychology, 13, 1010836.

38. Uddin, L. Q., Supekar, K., & Menon, V. (2013). Reconceptualizing functional brain connectivity in autism from a developmental perspective. Frontiers in Human Neuroscience, 7, 458.

39. Uljarević, M., Baranek, G., Vivanti, G., Hedley, D., Hudry, K., & Lane, A. (2017). Heterogeneity of sensory features in autism spectrum disorder: Challenges and perspectives for future research. Autism Research: Official Journal of the International Society for Autism Research, 10(5), 703–710.

40. Uscătescu, L. C., Kronbichler, M., Said-Yürekli, S., Kronbichler, L., Calhoun, V., Corbera, S., Bell, M., Pelphrey, K., Pearlson, G., & Assaf, M. (2023). Intrinsic neural timescales in autism spectrum disorder and schizophrenia. A replication and direct comparison study. Schizophrenia (Heidelberg, Germany), 9(1), 18.

41. Wang, X.-J. (2002). Probabilistic decision making by slow reverberation in cortical circuits. Neuron, 36(5), 955–968.

42. Watanabe, T., Rees, G., & Masuda, N. (2019). Atypical intrinsic neural timescale in autism. eLife, 8. 10.7554/eLife.42256

43. Wengler, K., Goldberg, A. T., Chahine, G., & Horga, G. (2020). Distinct hierarchical alterations of intrinsic neural timescales account for different manifestations of psychosis. eLife, 9. 10.7554/eLife.56151

44. Zhang, H., Schneider, T., Wheeler-Kingshott, C. A., & Alexander, D. C. (2012). NODDI: practical in vivo neurite orientation dispersion and density imaging of the human brain. NeuroImage, 61(4), 1000–1016.

